# Majority-LCL: Malaria Cell Detection via Label Contrastive Learning and Majority Voting Ensembling

**DOI:** 10.1101/2024.07.22.604640

**Authors:** Shreyan Kundu, Rahul Talukdar, Nirban Roy, Semanti Das, Soumili Basu, Souradeep Mukhopadhyay

## Abstract

Malaria is a contagious disease caused by Plasmodium, a group of single-celled parasites, and is most commonly transmitted by an infected female Anopheles mosquito. More than 40 percent of the global population is at risk, with approximately 219 million reported cases and around 435000 deaths recorded in 2017 alone. Despite the availability of several advanced diagnostic tools, accurate malaria diagnosis remains challenging in resource-constrained settings, where microscopists often struggle to improve diagnostic accuracy. Deep learning-based cell image classification helps reduce incorrect diagnostic conclusions by enabling automated analysis. This research aims to improve diagnostic accuracy by classifying malaria-infected cells using a majority voting ensemble framework combined with triplet loss aided label contrastive learning. Experimental results demonstrate the effectiveness of the proposed method on microscopic cell images in terms of accuracy, precision, recall, and other evaluation metrics.

## I. Introduction

THE Plasmodium group of single-celled protozoan parasites is the infectious agent that causes malaria. The disease is primarily spread through the bite of an infected female Anopheles mosquito. With around 240 million cases reported annually, the disease puts approximately 40% of the people at danger worldwide. Malaria primarily afflicted countries in Africa (Fig. 1). Two-thirds of malaria-related deaths worldwide occur in children under the age of five, accounting for 92% of malaria-related deaths in Africa. Common symptoms of malaria include fever, headaches, nausea, and in severe cases, seizures, jaundice, and coma, which can lead to death.

**Fig. 1.**
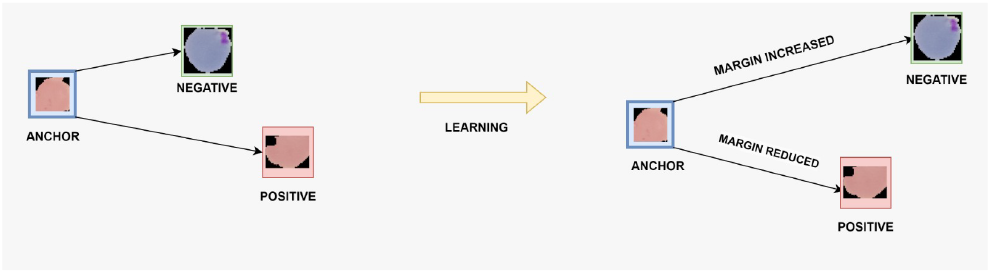
Visualization of triplet loss in our method

Every year, qualified professionals analyze millions of blood samples to look for signs of malaria infection. Malaria detection relies on manually counting parasites and infected red blood cells, a process heavily dependent on the microscopist’s skill and experience. Working in an environment with few resources and without a system that would aid in skill maintenance will impact diagnostic quality. This results in incorrect diagnostic choices.

## II. Related Work

The categorization of cell pictures using deep learning can help avoid making incorrect diagnosis conclusions. A type of machine learning known as deep learning has demonstrated impressive results across numerous non-medical fields. Deep learning has had little use in the medical industry because of privacy issues and a lack of specialized understanding in that area. However, deep learning has been applied in numerous medical domains recently. The paper by Liang et al. [1] was the first to use deep learning convolutional neural networks (CNNs) for malaria diagnosis.

A study by Odu et al. [2] built a machine learning model using XGBoost to predict malaria outbreaks in six African nations based on climate data. While successful, the model’s dependence on potentially limited climate data hinders realtime application.

Mahdieh Poostchi et al. [3] conducted a comprehensive review of image analysis and machine learning techniques for malaria detection from blood slides, discussing topics such as imaging, preprocessing, parasite detection, cell segmentation, and classification. However, the study lacks a consolidated framework, requiring substantial customization for practical implementation.

Narayanan [4] evaluated a range of machine learning and deep learning approaches for detecting malaria from cell images. They introduced a new Convolutional Neural Network (CNN) and contrasted it with established models like AlexNet, ResNet, VGG-16, and DenseNet. Despite achieving 96.7% accuracy and 0.994 AUC, computational demands restrict deployment in resource-constrained environments. Reddy et al. [5] implemented transfer learning using the ResNet-50 model to classify malaria-infected cells. The model showed high diagnostic accuracy, addressing challenges faced by microscopists in resource-constrained regions. However, one disadvantage is that transfer learning models like ResNet-50 require significant computational resources and may not be suitable for real-time applications in low-resource environments. A malaria evaluation system was created by Manning et al. [6] by employing image analysis and a feed-forward neural network (FNN) for cell classification. The system achieved 92% accuracy on a benchmark dataset, offering a promising tool for automated malaria diagnosis. Nonetheless, the primary limitation is its relatively lower accuracy compared to other advanced machine learning models, potentially reducing its reliability for clinical use. Ma et al. [7] study leverages active learning to improve malaria cell classification, achieving 93% accuracy using only 26% of the dataset. The research demonstrates that active learning, particularly uncertainty sampling, outperforms random sampling. However, a notable disadvantage is the increased computational time required for training. Related contrastive and ensemble learning strategies have also been explored in agricultural pest classification [17] and attention-driven medical image analysis [18], while swarm-based intelligent systems [21] indicate future opportunities for deploying such diagnostic frameworks in distributed and resource-constrained environments. Das et al. [8] employ machine learning to automate malaria parasite detection from microscopic images, achieving 84% accuracy. Their approach integrates feature selection using F-statistic, Bayesian learning, and SVM, highlighting complexities in optimizing classification performance due to the diverse nature of malaria stages and image variability.

This is how the rest of the paper is structured. Our motivation and contribution are illustrated in Section III. Our proposed method for classifying photographs of malaria cells is presented in Section IV. The trials we conducted and the results are detailed in Section V. This finding and the necessary tasks are outlined in Section VI.

## III. Motivation and Contribution

The reviewed studies underscore the significant advancements in using machine learning and deep learning techniques for malaria detection and prediction. While these methods demonstrate high accuracy and potential for improving malaria diagnosis, several limitations persist, such as reliance on highquality data, computational intensity, and challenges in real-world applicability. Addressing these drawbacks is crucial for developing robust and scalable solutions for malaria control and prevention.

The main contributions of this study are as follows:

1. To the best of our knowledge, the malaria cell classification problem is the first to use the combination of triplet loss (a contrastive environment) and cross-entropy loss (a supervised setting). By combining contrastive and supervised learning techniques for improved feature representation, this innovative method improves the model’s ability to categorize pests with more accuracy.
2. In contrastive learning, using a label memory bank reduces the number of erroneous negative examples. Through the retention and use of label data from prior batches, the model enhances its ability to discriminate between instances that are similar and those that are not, hence enhancing representation learning and overall performance by mitigating the influence of false negative samples.
3. Finally, the application of a majority voting-based ensembling of 3 top-performing label contrastive learning backbones (Resnet, Densenet and Efficientnet) proves the efficiency of our method in terms of accuracy. Recent studies have also explored contrastive and ensemble-based deep learning frameworks for medical image analysis beyond malaria detection. Kundu et al. [14] proposed a Gini-based feature selection and production-inspired feature fusion strategy for carcinoma grading, demonstrating the effectiveness of structured feature learning in histopathological images. In another work, Kar et al. [15] introduced an edge-guided TransEfficientUNET architecture for colon polyp segmentation, highlighting the importance of edge-aware representations in medical image segmentation tasks. Similarly, ensemble-based attention-driven models have been successfully applied to agricultural disease identification, as shown by Kundu et al. [16], indicating the robustness of ensemble learning in visual classification problems.

## IV. Our Method

### A. Label Contrastive Learning with triplet loss

Label Contrastive Learning was initially introduced by Yang et al. [10] for image classification using InfoNCE loss as contrastive loss. However, we have modified their approach slightly by substituting Triplet loss for InfoNCE loss. Supervised learning and contrastive learning are its two fundamental parts. The model distinguishes between various image classes through supervised learning. While the contrastive learning component pushes examples of different classes as far away as is practical, the supervised learning component encourages the model to position examples of the same class close to one another in the representation space. The contrastive learning component consists of two identical image encoders, *IE*_*q*_ and *IE*_*k*_. They are responsible for encoding the key image’s and input query picture’s representations, respectively. Encoder *IE*_*q*_ is updated via back-propagation, but encoder *IE*_*k*_ is updated via momentum. The label memory bank contains both the image representations and the corresponding class labels. The total loss of the model is the sum of the weighted classification loss and contrastive loss.

We treat the contrastive learning part of our design like a dictionary look-up problem. We query *Q* (a set of sample representations) for an encoding query, and the dictionary’s associated positive key, *Z*+. A contrastive loss requires a considerable loss between the query *Q* and the negative key *Z*−, as well as a little loss between the positive key *Z*+ and the query *Q*. To update the parameters of the *IE*_*k*_ encoder, the momentum update strategy proposed in MoCo [11] is employed, while the label memory bank method proposed in [12] is used. Below, we shall delve deeply into each element of the Contrastive loss formulation.

#### 1) Explanation of Image encoders

Two encoders, *IE*_*q*_ and *IE*_*k*_, are used to extract features from a cell image to create its feature representation. While *IE*_*k*_ pulls features from the key image, *IE*_*q*_ pulls features from the input query image. The model uses these extracted features for subsequent processing or comparison.

#### 2) memory bank with label (class)

Greater batch sizes in contrastive learning lead to higher learned representation quality. However, memory capacity limits the batch size, which cannot be expanded indefinitely. Ren et al. [12] proposed a label memory bank for our model, which is implemented with queues. Unlike the memory bank, the label(Class) memory bank saves both the related label and the image representation.

#### 3) Update on momentum

As the label memory bank contains a large number of picture representations and labels, back-propagation updating of the encoder *IE*_*k*_’s parameters can result in a significant overhead. The momentum update strategy reported in MoCo [11] can preserve image representations in the memory bank while minimizing computational cost. Furthermore, our method makes use of momentum update, which conducts gradient update operations for encoder *IE*_*q*_ solely by back-propagation. For updates to the encoder *IE*_*k*_ parameter, the momentum update technique is used; the precise update computation is as follows:

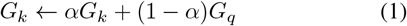

where *G*_*k*_ and *G*_*q*_ are the parameters of *IE*_*k*_ and *IE*_*q*_, respectively, and *α* is the momentum parameter, with possible values ranging from 0 to 1. According to MoCo, using a large momentum parameter *α* leads to the best results.

#### 4) Classification loss

The fully connected layers *IE*_*q*_ encoder output is sent to a classifier, which computes the classification loss. Here is the definition of a cross-entropy loss function, which is the classification loss function.

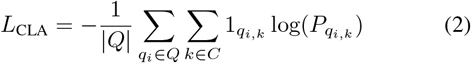

where *Q* is the set of representations for the query image, *P* is the estimated probability, 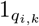 is the value of the indicator function, which is 1 when the query image’s classification is accurate, and *q*_*i,k*_ is the prediction that the query image *q*_*i*_ will be *k*.

#### 5) Triplet Loss as Contrastive loss

Let the i-th sample of our dataset *J* be denoted by (*I*_*i*_, *C*_*i*_). The feature representation of the picture *R*_*i*_ = *IE*(*I*_*i*_) is obtained by passing it through any image encoder *IE*. During the neural network training process, training samples are selected and generated into triplets. Each triplet 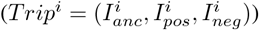consists of an anchor sample (*I*_*anc*_), a positive sample (*I*_*pos*_), and a negative sample (*I*_*neg*_). The tiplet labels are as follows: *C*_*anc*_ = *C*_*pos*_ ≠ *C*_*neg*_.Triplet loss aims to bring samples in the same class into neighboring locations on a manifold surface and push samples with different labels apart. The optimisation objective of the triplet *Trip*^*i*^ is,

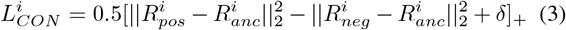

where,

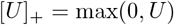

and δ is a hyperparameter that helps us to maintain 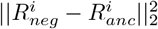 s to be greater than the distance 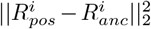. For all triplet *Trip*^*i*^ in the training dataset, the final objective to minimize is

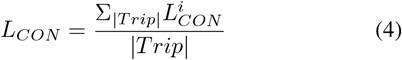

where *Trip* is the number of triplets in the training set. The visualization of triplet loss in our method is shown in Figure 1.

#### 6) Overall Loss

Our loss is the weighted sum of the classification and contrastive losses. Here’s how the overall loss is computed:

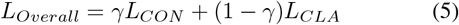

The hyper-parameter *γ* affects the relative weight of contrastive loss and classification loss. This method seeks to minimize *L*_*Overall*_, the contrastive loss that determines where in the representation space similar images are located as close together and other classes of images as far apart. The classification loss pushes the model to distinguish between many image classes. Once the complete model has been trained, pictures can be classified directly using the encoder *IE*_*q*_ and classifier. The triplet loss aided LCL method is shown in Figure 2.

**Fig. 2.**
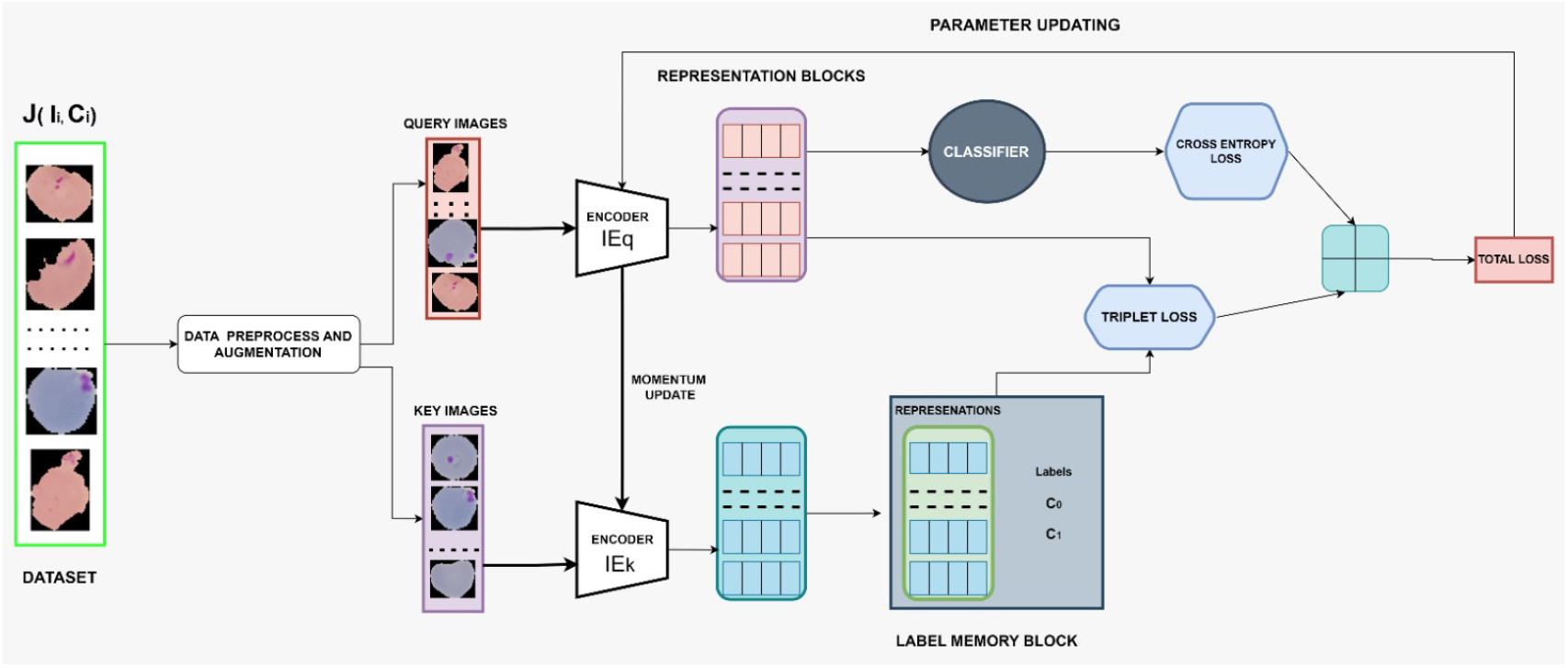
Triplet Loss aided LCL

### B. Majority voting based Ensembling

In our LCL-based architecture, we have shown that EfficientNet, Densenet, and Resnet are the top 3 performing backbones. So we apply majority voting to ensemble on the predictions of the 3 LCL-backbones to get more efficiency in the malaria cell image classification problem. Our overall method is shown in Figure 3.

**Fig. 3.**
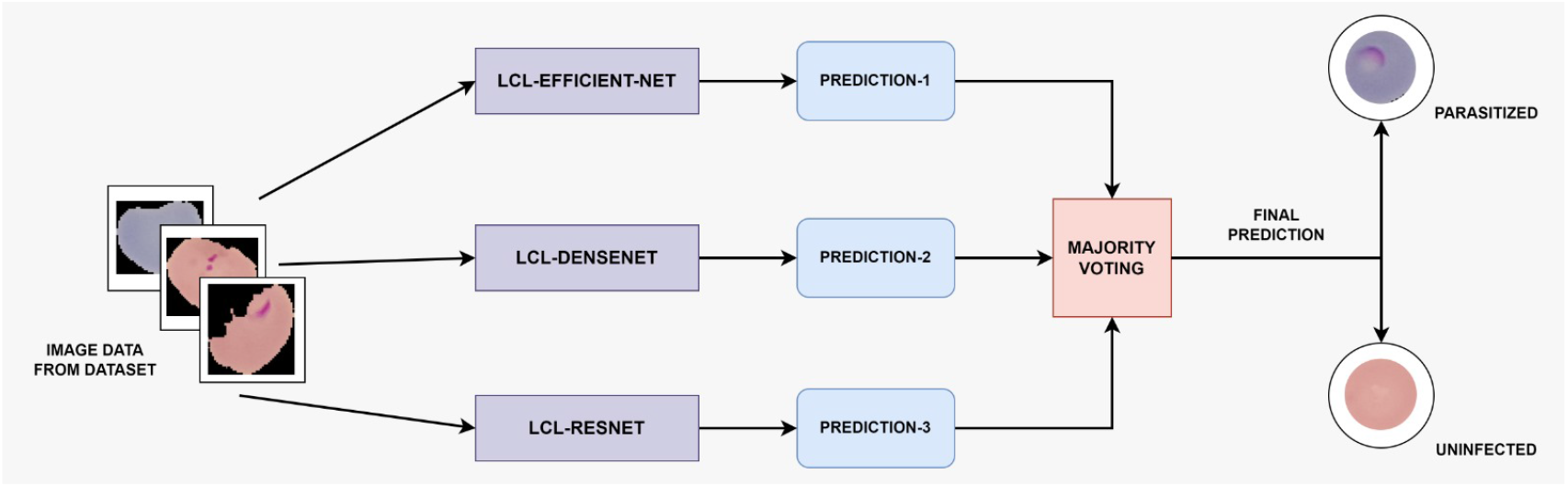
Majority Voting Based LCL-Ensembling

## V. Results

### A. Dataset Description

The NIH website is the source of this collection [9]. The images were meticulously identified by a professional slide reader at the Mahidol-Oxford Tropical Medicine Research Unit in Bangkok, Thailand. The de-identified images and annotations are kept at NLM. The collection consists of 27,558 cell images, with equal numbers of parasitised and uninfected cells. In contrast to negative samples, which lacked impurities or staining artefacts, positive samples contained Plasmodium. A train set (70%), a validation set (20%), and a test set (10%) comprise our dataset.

### B. Experimental setup

The computer used for all of the trials has a 12-GB NVIDIA Tesla T4 GPU, and Python 3.6 is the programming language used. The deep learning model is created in the Tensorflow environment using the Keras library.

### C. Parameters and hyperparameters

Our approach carefully chooses its parameters and hyperparameters to maximise performance. We use the Stochastic Gradient Descent (SGD) optimiser with a weight decay of 10^−4^ and a batch size of 32. To balance the loss components, we set *α* to 0.9 and set the temperature hyperparameter to 0.5. The model is trained across ten epochs with a fixed learning rate of 0.001. In order to manage the contributions of different terms in the loss function and guarantee optimal convergence during training, we also set δ to 0.5 and *γ* to 0.3.

### D. Evaluation parameters

A test gets results in 4 ways: correctly identified positive (True Positive-TP), correctly identified negative (True Negative-TN), mistakenly called negative (False Negative-FN) and mistakenly called positive (False Positive-FP). We assess our model using established measures such as F measure, Accuracy, Precision, and Recall.

### E. Our Result and visualization

We have got 97.53% as overall accuracy in the test dataset. Similarly, we got 97.4% as precision, recall, and F-score. Here, class 0 means malaria infected cell and class 1 means uninfected cell. For a better understanding of the readers, we have computed the confusion matrix as well as the ROC curve in Figure 4. The AUC score is 0.99. Now, we have shown TSNE to prove the efficacy of our classification algorithm in Figure 5.

**Fig. 4.**
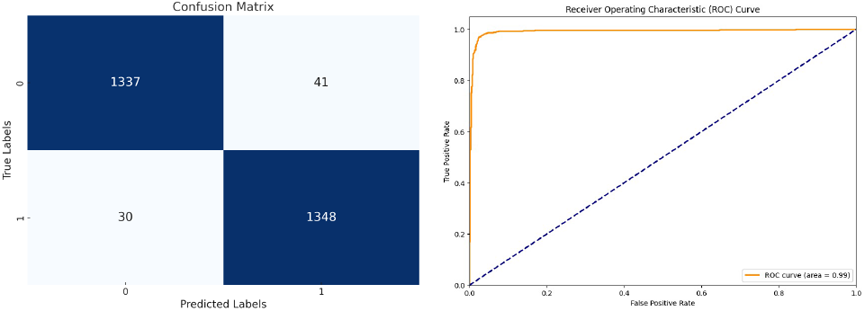
Confusion matrix and Roc curve

**Fig. 5.**
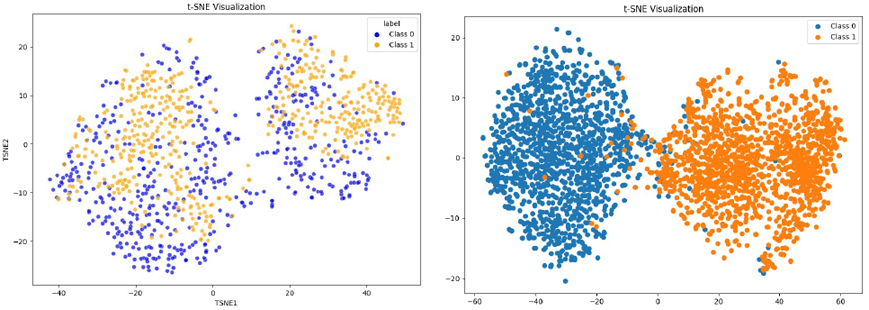
TSNE before and after our method

### F. Comparison with different backbones

The subsection presents a numerical comparison of our method with different CNN backbones, both with and without LCL. Without LCL, our method achieves the highest accuracy (97.53%) compared to EfficientNetB3 (97%), DenseNet121 (96.23%), and MobileNetv3 (96.02%). For precision, recall, and F1 score, our method achieves 97.4%, outperforming all others. With LCL, our method still shows superior accuracy, ahead of DenseNet (97.31%) and ResNet50 (97.02%). Precision, recall, and F1 score for our method remain the highest (97.4%), demonstrating its consistent superiority in both LCL and non-LCL settings across all metrics. Figure. 6 represents the graphical comparisons.

**Fig. 6.**
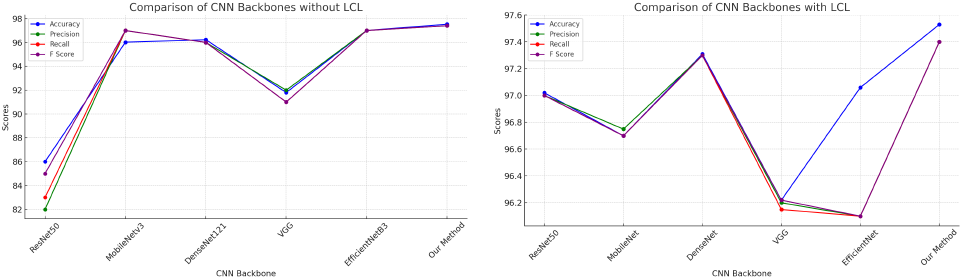
Comparison of different CNN backbones with our method

### G. Comparison with SOTA

Table I compares our method with state-of-the-art (SOTA) approaches from the literature. Our method achieves the highest accuracy (97.53%) compared to Liang et al. [1] (91.99%) and Manning et al. [6] (92.2%), demonstrating superior performance. In terms of precision, recall, and F1 score, our method also outperforms others, with all three metrics reaching 97.4%. In contrast, Liang et al. [1] achieve lower precision (95.12%) and recall (89%), while Das et al. [8] achieve high recall (96.62%) but lower accuracy (83.5%). Moreover, while some methods perform well in certain metrics, such as high recall or precision, they fail to demonstrate a balanced performance across all indicators. Our method not only excels in accuracy but also provides robustness across precision, recall, and F1 score, making it a comprehensive solution compared to these SOTA approaches.

**TABLE I.** Comparison of Our Method with different SOTA.

| Backbone | Accuracy | Precision | Recall | F score |
| --- | --- | --- | --- | --- |
| Liang et al. [1] | 91.99 | 95.12 | 89 | 90.24 |
| Reddy et al. [5] | 95.4 | - | - | - |
| Das et al. [8] | 83.5 | 88.51 | 96.62 | 92.38 |
| Ma et al. [7] | 75 | - | - | - |
| Manning et al. [6] | 92.2 | - | - | - |
| **Our Method** | **97.53** | **97.4** | **97.4** | **97.4** |

## VI. Takeaways

Our research created a new label contrastive learning architecture with triplet loss and majority voting ensembling method, testing several backbones and selecting Resnet, Densenet and Efficientnet for its high accuracy (97.53%). This label contrastive approach, combining triplet loss and crossentropy, achieved excellent precision, recall, and F-score. We believe this approach, along with careful hyperparameter tuning, paves the way for future advancements in label contrastive learning. Future work exploring dual backbones, and various ensembling techniques in broader applications in medical imaging and other areas is planned. This research lays a strong foundation for further exploration in this rapidly evolving field. Future extensions of this work will explore game-theoretic and equilibrium-based formulations for segmentation and classification, inspired by recent advances in strategic segmentation frameworks [22]. Additionally, integrating uncertainty-aware ensembling and confidence-weighted decision mechanisms, as investigated in renal carcinoma grading [20], can further improve robustness in clinical deployment. The applicability of the proposed contrastive-ensemble paradigm will also be expanded to cross-domain medical and agricultural datasets, building upon prior studies in crop disorder detection and precision agriculture [13], [19].

